# Evolution of dispersal and the maintenance of fragmented metapopulations

**DOI:** 10.1101/2022.06.08.495260

**Authors:** Basile Finand, Thibaud Monnin, Nicolas Loeuille

## Abstract

Because it affects dispersal risk and modifies competition levels, habitat fragmentation directly constrains dispersal evolution. When dispersal is traded-off against competitive ability, increased fragmentation is often expected to select higher dispersal. Such evolutionary effects could favor the maintenance of the metapopulation by fostering spatial rescue effects. Using an evolutionary model, we first investigate how dispersal evolves in a metapopulation when fragmentation and aggregation of this fragmentation are fixed. Our results suggest that high fragmentation indeed selects for dispersal increase, but this effect is largely reduced in aggregated landscapes, to the point of being nonexistent at the highest aggregation levels. Contrasted dispersal strategies coexist at high fragmentation levels and with no or low aggregation. We then simulate time-varying fragmentation scenarios to investigate the conditions under which evolutionary rescue of the metapopulation happens. Faster evolution of dispersal favors the persistence of the metapopulation, but this effect is very reduced in aggregated landscapes. Overall, our results highlight how the speed of evolution of dispersal and the structuration of the fragmentation will largely constrain metapopulation survival in changing environments.

## INTRODUCTION

Dispersal, defined as the movement of individuals associated with gene flows across space (Ronce, 2007), is a key process in ecology and evolution. It has important consequences for population dynamics, changes in species distribution, maintenance of genetic diversity and for local adaptation (Travis et al., 2013). Habitat loss and fragmentation result in decreased population sizes and gene flows, which undermines population viability and ultimately species survival. In landscapes that include suitable and unfavorable patches of varied size and distribution, dispersal allows individuals to move between suitable patches thereby favoring the survival of the metapopulation through spatial rescue effects (Levins, 1969). In a source-sink context, dispersal increases spatial occupancy as source populations allow the persistence of peripheral sink populations through dispersal (Pulliam, 1988). The maintenance of sink populations is especially important in the context of current changes as source-sink hierarchies could change in time. Given such environmental changes, dispersal helps the survival of species by allowing them to follow suitable niche conditions, thereby playing a key role in range expansions (Phillips et al., 2006).

Habitat fragmentation creates spatial heterogeneities in several ways. It decreases the quantity of suitable habitat by decreasing the size and increasing the isolation of suitable patches, even though it increases their number (Fahrig, 2003). In our study, fragmentation is defined by the proportion of hostile vs. suitable locations (patches) and we systematically vary its degree of spatial aggregation. Dispersal evolution is affected by fragmentation, due to variations of different selective pressures. By definition, fragmentation increases spatial heterogeneity so that dispersing propagules encounter non-suitable patches more frequently. Theoretical and empirical studies suggest that such increases in dispersal costs and in spatial heterogeneity select decreased dispersal (Bonte et al., 2006; Cheptou et al., 2008; Duputié & Massol, 2013; Hastings, 1983; Schtickzelle et al., 2006; Travis & Dytham, 1999). While such a counterselection of dispersal was originally highlighted in theoretical models (Hastings, 1983; Travis & Dytham, 1999), empirical evidence for such effects has accumulated in recent years, for a large variety of species, from the weed *Crepis sancta* (Cheptou et al., 2008), to the butterfly *Proclossiana eunomia* (Schtickzelle et al., 2006) and the wolf spider *Pardosa monticola* (Bonte et al., 2006). Habitat fragmentation however also increases inbreeding, kin competition or temporal variation of the environment and all of these components usually select for higher dispersal abilities (Charlesworth & Charlesworth, 1987; Cote et al., 2017; Duputié & Massol, 2013; Gandon, 1999; Hamilton & May, 1977; Matthysen et al., 1995; Oldfather et al., 2021; Tung et al., 2018). In addition to the modulation of overall dispersal levels, fragmentation can also, under certain conditions, maintain contrasted dispersal strategies simultaneously. Previous investigations suggest that such a dispersal polymorphism evolves under high fragmentation and high aggregation, with dispersing and non-dispersing individuals coexisting within the same population (Bonte et al., 2010). It principally appears because aggregation produces a coexistence of many small patches and few large patches (Massol et al., 2011; Parvinen, 2002; Parvinen et al., 2020), or due to edge effects that select low dispersers at the edge and high dispersers in central places (Travis & Dytham, 1999).

While these previous studies consider dispersal as an isolated trait, it is now widely recognized that evolutionary changes in dispersal most often imply variations in phenotypic traits that constrain other ecological interactions (Raffard et al., 2022). It has been highlighted that when colonization abilities (here our measure of dispersal) are traded against competitive abilities, coexistence of a large number of strategies is possible along this hierarchy (Tilman, 1994). This trade-off has a long history in ecology and former studies investigated how it may explain the coexistence of species within metacommunities (Calcagno et al., 2006; Tilman et al., 1997; Yu & Wilson, 2001) and how such a diversity varies when fragmentation or habitat destruction occurs (Tilman et al., 1994, 1997). While these studies mostly focused on ecological dynamics, we here use the trade-off to investigate its eco-evolutionary implications in a fragmentation context. Such a trade-off could for instance occur because given a fixed quantity of energy, allocation could produce a large number of small propagules (colonizer) or few large propagules (competitor) (eg, Geritz et al., 1999; Smith & Fretwell, 1974). For example, the weed *Crepis sancta* produces small and/or large seeds. Small seeds have high wind dispersal due to their lightweight but low competitiveness due to low resource storage. In contrast, large seeds have restricted dispersal due to their weight but contain more resources (Cheptou et al., 2008). In social insects, dispersal and reproduction could follow from the production of many isolated queens that fly large distances and have high mortality or through the split of the colony in a few propagules that usually disperse on short distances but may be more efficient at gathering resources when founding the new colonies (Cronin et al., 2013, 2016). Habitat fragmentation affects strategies along the competition-colonization trade-off in different ways. First, it directly lowers the average density at the metapopulation level, thereby changing competitive pressures. Second, it creates isolated patches that act as a positive filter for the best dispersers. To our knowledge, only one study considers how this competition/colonization trade-off affects the dispersal strategies selected by fragmentation (Tilman et al., 1994). This study shows that in a spatially variable environment with an increase of fragmentation, the more competitive (and thus the less dispersive) strategies disappear first, so that high dispersal strategies are selected.

Such results are obtained without considering explicit spatial structures as the position of patches is not accounted for in Tilman et al. (1994) (mean field approximation). Fragmentation of the environment can however be an aggregated process, as human activities such as urban development or agricultural exploitation are often concentrated in specific locations. A previous work on metapopulations shows that the structuration of habitat heterogeneities is crucial to study metapopulation responses to fragmentation (Hiebeler, 2000).

When environmental heterogeneities are spatially correlated (aggregation), predictions based on mean-field approximation are often qualitatively incorrect when compared to spatially explicit approaches (Hiebeler, 2000). In contrast, mean-field approximations yield correct results in the case of randomly distributed fragmentation. Leaving out the competition/colonization trade-off, the importance of aggregation in the evolution of dispersal is highlighted by various studies (Bonte et al., 2010; Fronhofer et al., 2014; Ovaskainen et al., 2002; Travis & Dytham, 1999). For example, in the context of correlated extinctions, empirical work on the spider mite *Tetranychus urticae* and an associated theoretical model show a selection for long-distance dispersal and a decrease of local dispersal compared to spatially random extinctions (Fronhofer et al., 2014). Travis and Dytham (1999) found a decrease in dispersal with increased fragmentation, but an increase in dispersal with higher aggregation. The risk to disperse outside of a large aggregate of suitable patches and into a hostile environment is indeed lowered, so that aggregation modulates dispersal costs. Similarly, Bonte et al. (2010) found a decrease of local and global dispersals with the increase of fragmentation, and demonstrates that decreasing aggregation has the contrasted effect of decreasing local dispersal and increasing global dispersal. To summarize, the study that considers variations of dispersal strategies along a competition/colonization trade-off in fragmented habitats use a spatially implicit (mean field) approach, while others use spatially explicit landscapes but ignore possible competition/colonization trade-offs. The goal of our study is therefore to integrate both aspects, that is to study the evolution of dispersal along the competition/colonization trade-off given a spatially explicit structuration of the habitat.

Understanding this dispersal evolution has immediate consequences to better predict the maintenance of metapopulations. For instance, a selected increase in dispersal favors the exchange of individuals between patches and the colonization of empty patches (spatial rescue). Extinction may also be prevented, by the emergence of evolutionary rescue, when natural selection favors adapted traits (Bell, 2017; Carlson et al., 2014; Gomulkiewicz & Holt, 1995). Here, an evolutionary increase of dispersal distances could avoid a population extinction in a climate change context (Boeye et al., 2012) or in a context of high mortality (Heino & Hanski, 2001). Given a temporally increasing fragmentation, natural selection may favor high dispersal, as the availability of empty and isolated patches constantly increases. Because only highly dispersive strategies can reach them, such isolated patches act as filters that favor high dispersal (Heino and Hanski, 2001). Consistent with this theoretical prediction, a temporal increase of fragmentation led to higher dispersal in *Drosophila melanogaster* experiments (Tung et al., 2018). Conversely, if evolution were to lead to less dispersal, it would potentially decrease metapopulation persistence (Gyllenberg et al., 2002). The implication of the evolution of dispersal for metapopulation persistence in a world that becomes increasingly fragmented is therefore an important, unresolved issue.

Using metapopulation simulations, we studied how the spatio-temporal structuration of fragmented environments acts on dispersal evolution given a competition/colonization trade-off. First, we fixed fragmentation and aggregation levels and investigated how dispersal evolved. Second, we varied fragmentation over time to test whether dispersal evolution can prevent extinction (evolutionary rescue), under various rates of evolution of dispersal. We hypothesize that, in a fixed environment, higher fragmentation selects for an increase in dispersal because more empty patches will become available to colonizers and inaccessible to competitors. In addition, competition could be relaxed in fragmented landscapes as the average occupancy is lowered. However, if the fragmentation is aggregated, large groups of suitable patches could persist in the landscape. Such a situation is favorable to competitors and should decrease the selection toward higher dispersal or lead to dispersal polymorphism with competitors dominating aggregated patches while colonizers remain favored in isolated patches. When fragmentation increases over time, we hypothesize that the occurrence of evolutionary rescue depends on the speed of evolution of dispersal, which needs to be faster than the speed of fragmentation to counterbalance its effects.

## MODEL PRESENTATION

Simulations and analysis were done with R 3.9. Our simulations consider a spatially explicit environment consisting of a grid of 50×50 patches wrapped into a torus to avoid edge effects (Fig. 1). Each patch can be in one of three possible states: unsuitable, suitable and empty or suitable and occupied. Only suitable and empty patches are available to dispersing individuals. Importantly, we define fragmentation as the percentage of unsuitable patches. This definition of fragmentation is classically considered in the literature and is directly linked to other components often used to describe fragmentation such as the number of independent patches, their size or their isolation (Fahrig, 2003). For a given level of fragmentation, we independently vary the degree of aggregation of unsuitable patches, as controlled by the Hurst coefficient. This coefficient is directly related to how similarity among patches decrease with distance thereby constraining spatial autocorrelation. While we keep a simple definition of fragmentation (proportion of unsuitable patches), note (1) that higher frequency of unsuitable patches decreases overall connectivity; (2) that we also manipulate the effect of fragmentation on local contexts by considering varying degrees of aggregation. Examples of landscapes can be found in the upper left part of Fig. 1. Unsuitable patches are distributed randomly or with a set percentage of aggregation (created with a fractal Brownian motion) using the *NLMR* and *landscapetools* package (Sciaini et al., 2018). A higher aggregation means that a suitable patch is more likely to be close to another one compared to the random expectation.

**Figure 1:**
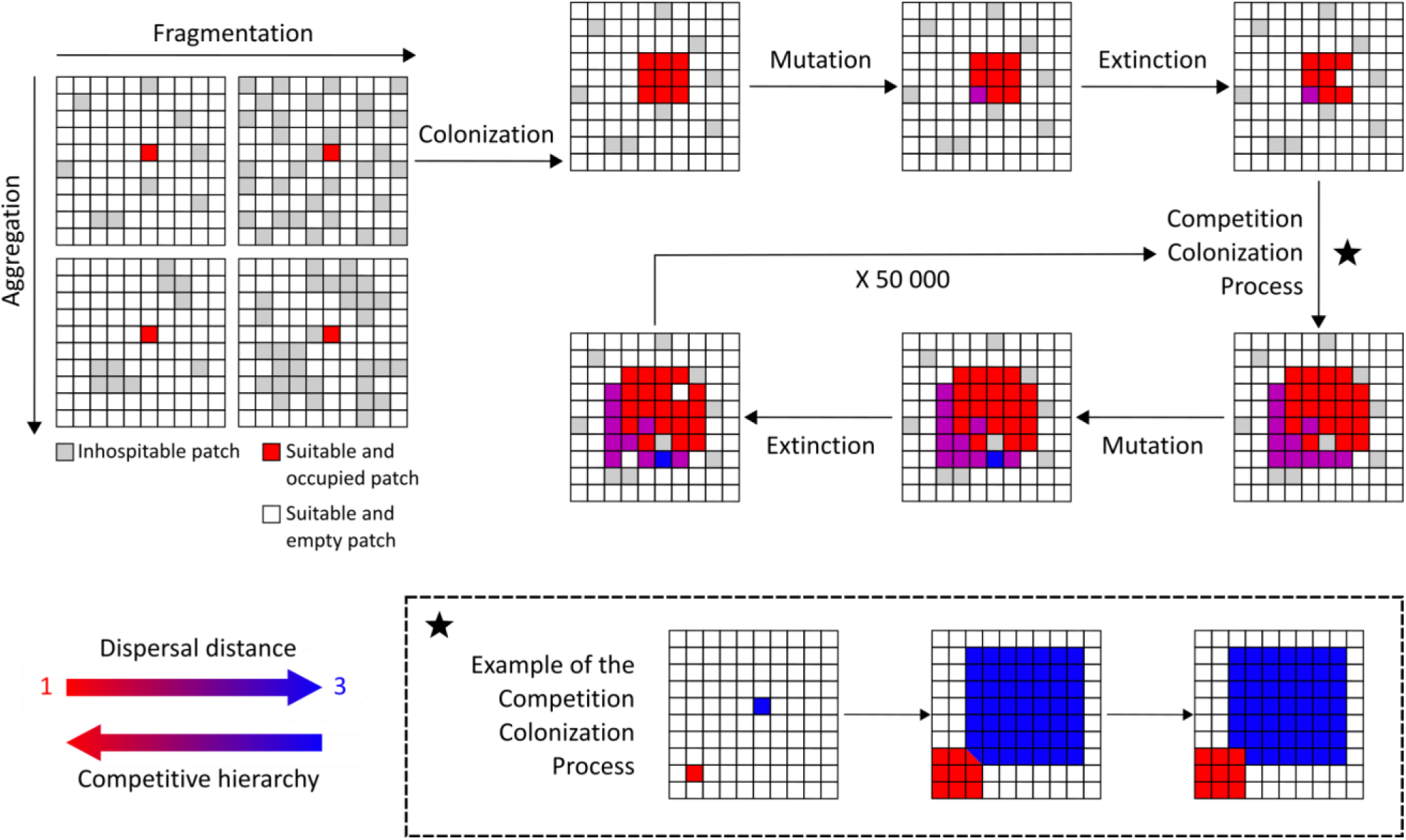
Illustration of a simulation based on the competition/colonization tradeoff in a fixed environment scenario. Upper panels detail the various parts of a given timestep, while the box below shows the competition/colonization process when two individuals arrive on the same patch. For each time-step, individuals colonize all suitable and empty patches within their dispersal distance. Individuals mutate with a small probability. If a mutation occurs, the dispersal distance of the individual is modified by 1, upward or downward with equal probability. Extinction events follow, with a probability *e*. If two (or more) individuals arrive on the same patch, the one with the smallest dispersal distance, being competitively dominant, wins the patch. One simulation lasts 50 000 time-steps.

Individuals are characterized by two traits: colonization and competition capacities (both integer values) directly linked through the colonization/competition trade-off. The model relies on discrete time steps, each time step being divided in three parts: colonization, competition and extinction (see Fig. 1).

1. Colonization. The colonization capacity defines the radius (number of patches) of the area around the individual where its offspring are dispersed. We assume that offspring will potentially colonize all empty but suitable patches within this range. This implicitly assumes that individuals with larger colonization capacity are not limited by the number of offspring they produce, assuming an increase of fecundity when dispersal distance increases (see introduction).
2. Competition. We assume that only empty suitable patches can be colonized by propagules. Given that individuals fill all suitable surrounding patches with their offspring, suitable empty patches are often reached by several offspring simultaneously. We then assume that the competitive hierarchy favors the strategy that has the smallest dispersal distance (competition-colonization trade-off, lower part of Fig. 1). The surviving individual inherits the dispersal strategy of its parents, except in the case of rare mutation events. When mutation occurs, the dispersal range of the mutant individual is enhanced or reduced by one cell, with equal probability. Mutations incur incremental variations in dispersal distance of 1, upward or downward, with equal probability. Dispersal distances below 0 are not possible and discarded. Note that while this situation is never observed here, a dispersal distance above 25 would mean global dispersal over the whole grid. We assume that established individuals (ie, occupied patches) cannot be displaced by incoming propagules, regardless of their traits.
3. Extinction. We assume that extinction probability does not depend on the dispersal trait.

Occupied suitable patches therefore become empty (but still suitable) with a fixed probability *e* at each time step (*e*=0.05).

Each landscape is populated, at the beginning of the simulation, with ten populations that are randomly distributed on suitable patches. These populations are assumed to be highly dispersive (colonization capacity of twelve). We verified that initial conditions (number of populations and initial colonization capacity) do not affect the equilibrium outcome (Supplementary information Figure 1).

### Scenario 1: Evolution of dispersal in fixed fragmented landscapes

In the first scenario, we fix the landscape and study how dispersal evolution depends on the levels of fragmentation and aggregation. Fragmentation corresponds to a specified percentage of unsuitable patches (i.e. 0, 20, 40, 60, 80, 90, 95 or 99% of patches are assumed unsuitable). These unsuitable patches are aggregated at varying degrees (0, 20, 40, 60, or 80%). To assess repeatability, twenty different landscapes are generated for each combination of fragmentation and aggregation. The mutation rate is set at 0.1. Each simulation lasts 50 000- time steps. Because simulations are stochastic, they never reach a completely stable equilibrium, but we visually checked for each simulation that 50 000 time steps allowed the system to reach a stable regime that can be characterized (Figure S2). It means that the mean and the variance stay stable over at least 5 000 time steps (more discussion is provided in supplementary information S2). We record the mean dispersal capacities of individuals during the simulation and the occupancy of each dispersal strategy.

### Scenario 2: Evolutionary rescue under progressively increasing fragmentation

In the second scenario, we progressively increase the level of fragmentation over time. We systematically manipulate the rates of fragmentation and of mutation to investigate conditions under which dispersal evolution can delay extinction. The grid is supposed to be fully suitable at the onset of the simulation and for the first 200-time steps. We then progressively increase fragmentation until the metapopulation becomes extinct. As in the first scenario, the increase in fragmentation occurs with random or aggregated distributions of unsuitable patches (levels of aggregation: 0%, 10%, 20%, 40%, 60%, 80%). Rates of fragmentation correspond to the probability that a suitable patch becomes unsuitable within a given time step. We tested three rates of fragmentation (0.0001, 0.001, 0.01). As evolutionary rescue is construed as a race between the speed of the disturbance and the speed of adaptation (Gomulkiewicz & Holt, 1995), we also systematically manipulate the speed of evolution by considering different rates of mutation (0.001, 0.01, 0.1). We replicate each combination of aggregation, fragmentation rate and mutation rate forty times. We record the fragmentation at population extinction as an index of the resistance of the metapopulation to the disturbance. Higher values of this index show that evolution of dispersal allowed the metapopulation to survive higher levels of the disturbance. Evolutionary rescue occurs if metapopulations with evolution of dispersal resist higher disturbance levels than metapopulations without dispersal evolution. For each set of simulations, we also record the variations of dispersal strategies (occupancy of the various dispersal phenotypes) over time to identify the path that evolutionary rescue takes.

## RESULTS

### Evolution of dispersal in fixed fragmented landscapes

Higher fragmentation selects for increasing mean dispersal distances. In non-fragmented landscapes, competitive strategies eventually dominate so that dispersal distance quickly evolves close to one (Fig. *2*a and b). Such a strategy remains dominant for all low levels of fragmentation (0 to 60%). High dispersal is selected under higher fragmentation, especially strongly when fragmentation is random (7.27±1.08 patches at 99%, red line in Fig. 2a, see also Fig. 2c). However, adding aggregation strongly decreases this selection effect. For instance, a little bit of aggregation (20%, orange line in Fig. 2a, see also Fig. 2d) suffices to lower the selected dispersal distance in very fragmented landscapes to 2.09±1.02 patches. Higher aggregation (40 to 80%) further decreases the selected dispersal distance, so that fragmentation hardly has any effect on selected dispersal when aggregation is high (blue lines, Fig. 2a). Aggregation therefore qualitatively changes the results of mean field models (such as Tilman et al., 1994).

**Figure 2:**
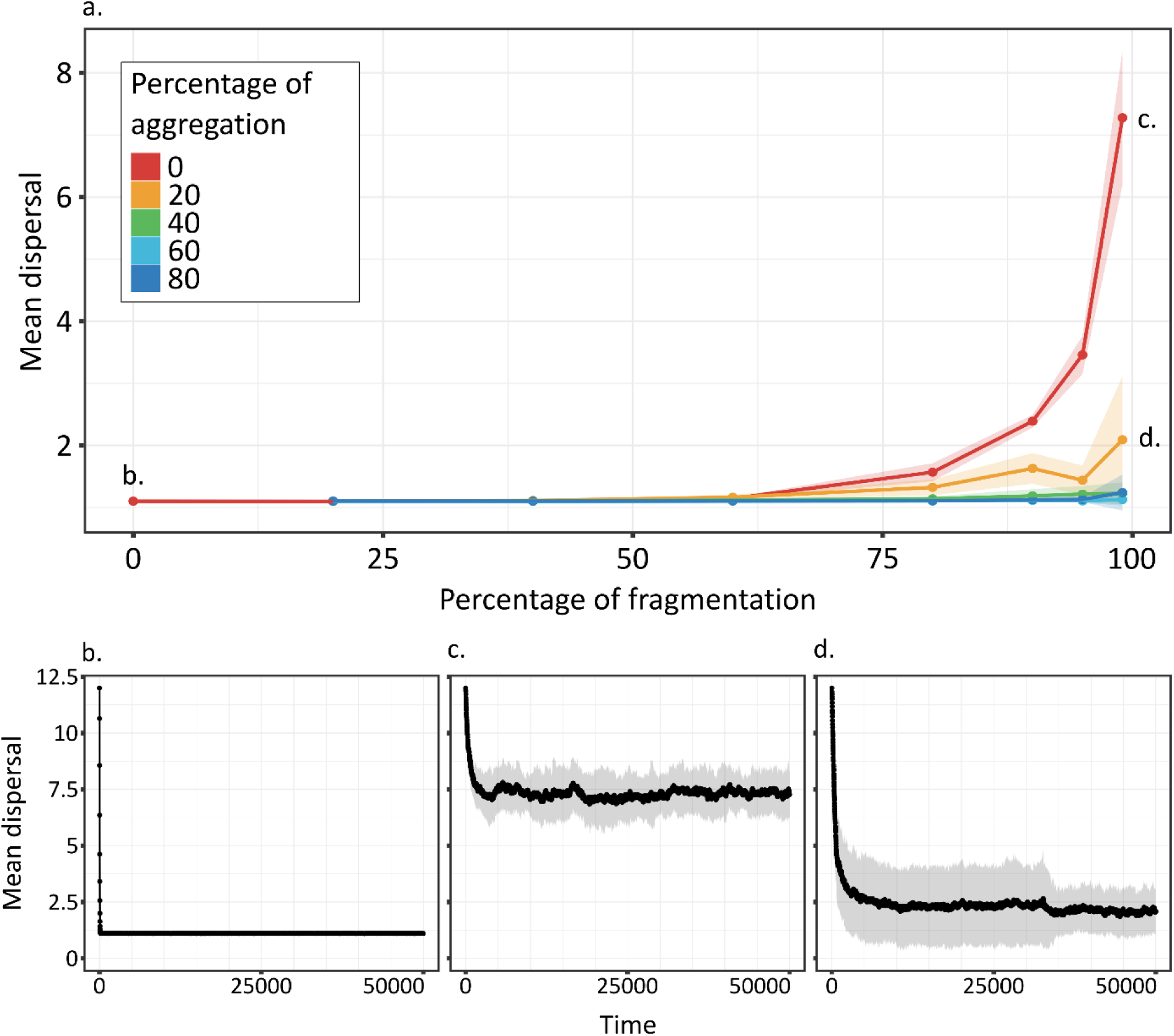
Dispersal (mean +/- SD) at the end of simulations (at equilibrium) depending on environment fragmentation and aggregation (a), and over time (b-d) for the 20 replicates for conditions of no fragmentation and no aggregation (b), 99% of fragmentation and no aggregation (c) or 99% of fragmentation and 20% of aggregation (d). Note that higher dispersal distance is selected in random fragmented landscapes, but that aggregation of fragmentation lowers this selective effect. Shadows around curves represent SD.

### Evolution of polymorphism in fragmented landscapes

Beyond the observed variations in mean dispersal distances, the long-term variability of dispersal strategies also depends on fragmentation and aggregation. Particularly, when fragmentation is sufficient, we observe the coexistence of several dispersal strategies (polymorphism, Fig. 3). In all cases of polymorphisms, we observe similar patterns. Suitable patches that are close to one another sustain the less dispersive strategies, while isolated patches act as filters that favor the more dispersive strategies. The set of polymorphic strategies however vary depending on the relative positions of patches. For instance, given a very high fragmentation (99%) with no or little aggregation (20% or less), few suitable patches are close to one another by chance (purple patches, Fig. 3b). Because the distances among these patches is still quite important, the dispersal strategy they sustain is still quite high (around 5). Other patches are more isolated (blue patches, Fig. 3c) and act as a selective pressure favoring very high dispersal distances (around 9). When fragmentation is slightly lower (80 to 95%) and aggregation slightly higher (20 to 40%), large aggregates of suitable patches occur in the landscape (red patches, Fig. 3e) and favor competitive strategies (dispersal distance around 1). The remaining suitable patches are isolated and favor a continuum of more dispersive strategies.

**Figure 3:**
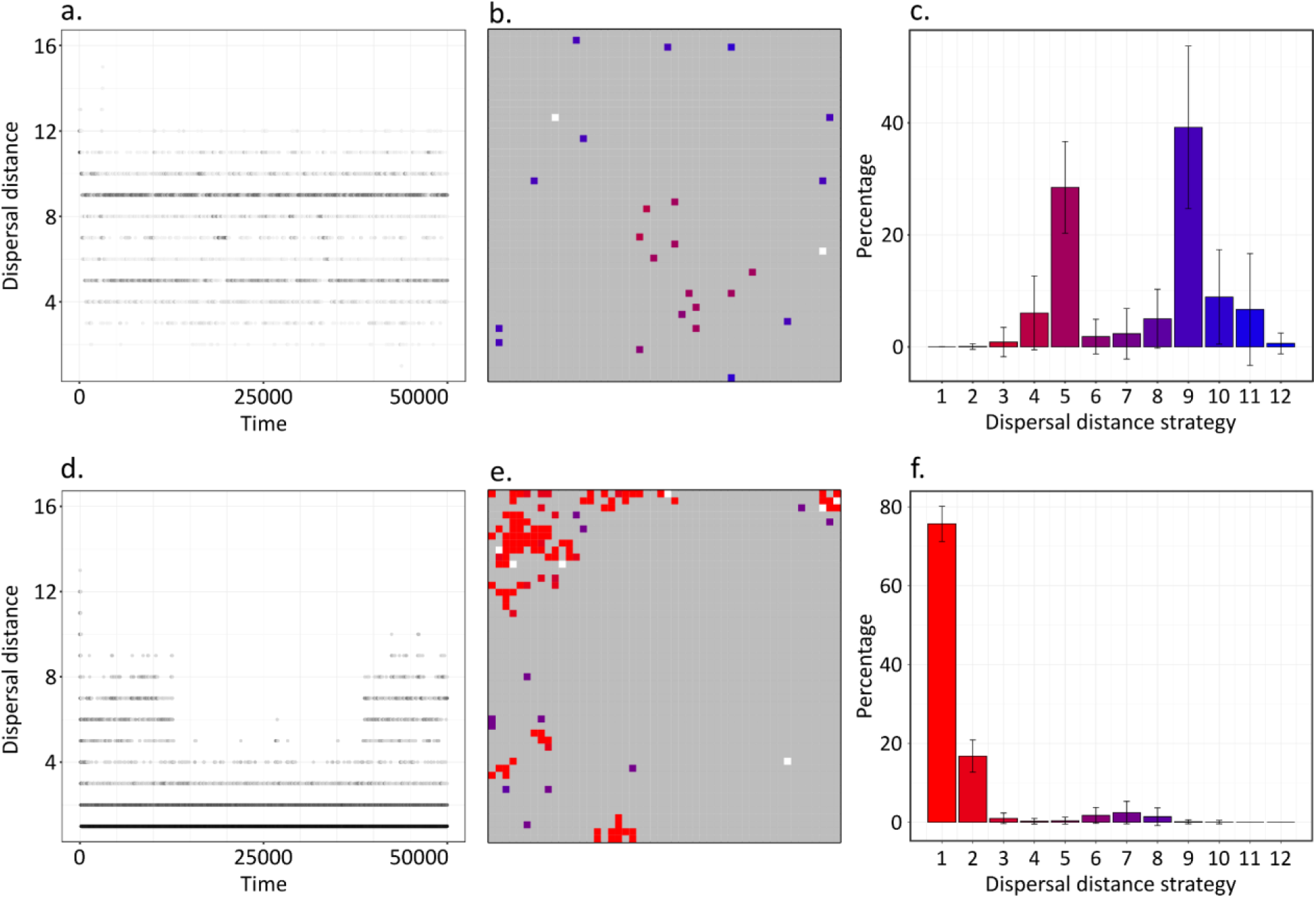
2 examples of simulations showing dispersal polymorphism. (a) and (d) show the presence of dispersal strategies over time in two separate simulations. (b) and (e) represent the grid at the end of the corresponding simulations. (c) and (f) show the relative abundance of each strategy (mean percentage +/- SD) over the last 5 000-time steps. For (a) and (d) the intensity of black represents the proportion of each strategy for the given time. It is log-transformed for (d). For (b) and (e) grey patches are unsuitable, white patches are suitable and empty, and coloured patches are suitable and occupied by populations differing in dispersal strategies (ranging from low dispersal in red to high dispersal in blue). Dispersal strategies are similarly color coded in panels (c) and (f). The first row of panels (a-c) shows an example with two equally abundant dispersal distance strategies (dispersing at 5 and 9 patches). Conditions are fragmentation of 99% and no aggregation. The second row of panels (d-f) shows an example where one dispersal distance strategy (at 1.1 patches) dominates (fragmentation of 95% and aggregation of 20%).

### Evolutionary rescue under progressively increasing fragmentation

When fragmentation increases over time, fast dispersal evolution allows a longer persistence of the metapopulation, i.e. an evolutionary rescue. Fig. 4 shows this evolutionary rescue as the difference (orange arrows) between the scenario with no evolution (mutation rate equal zero) and the three scenarios with more or less rapid evolution. Intuitively, evolutionary rescue occurs and is strongest when there is no aggregation, fragmentation rate is low, and mutation rate is high (mean difference of 3.04% between scenarios without and with evolution, Fig. 4a). Evolutionary rescue is largely decreased when fragmentation rate is higher (a difference of 0.63%, Fig. 4c). Variations in the potential of evolutionary rescue are not continuous. Rather, a jump in the extinction time when mutation rates increase can be identified. This jump is relative to the fragmentation rate. Under our set of parameter values, evolutionary rescue occurs when the mutation rate is ten times higher than the fragmentation rate (Fig. 4a,b,c, blue arrows). Finally, we stress that aggregation largely constrains evolutionary rescue. No potential for evolutionary rescue can be identified in aggregated landscapes (Fig. 4d,e,f).

**Figure 4:**
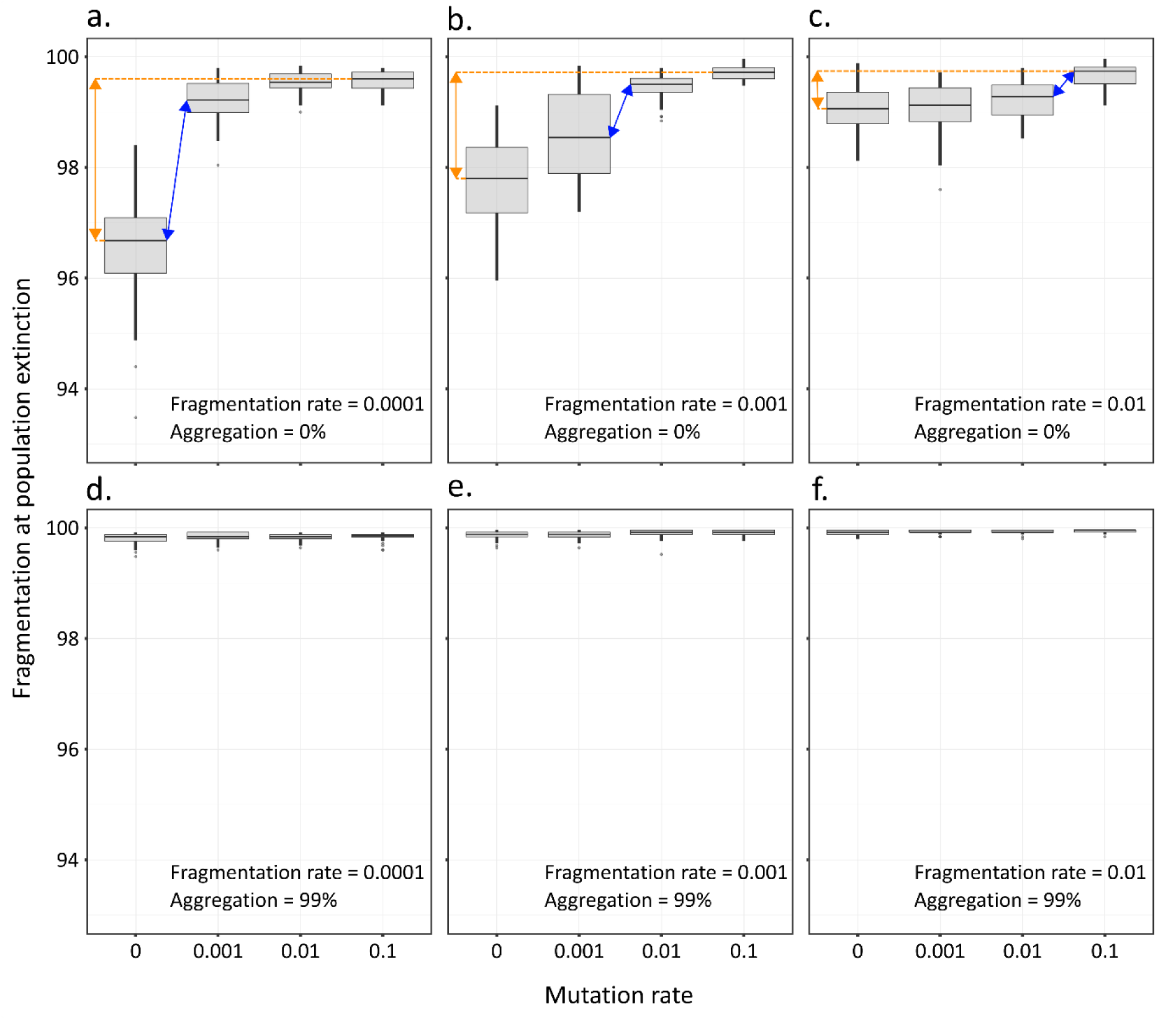
Fragmentation level leading to the metapopulation extinction versus mutation rate for different scenarios of fragmentation rate and aggregation. Orange arrows highlight the difference of extinction thresholds between no evolution and fast evolution, a proxy for maximal evolutionary rescue. Blue arrows highlight the change in mutation rate that has a maximal effect on evolutionary rescue. Note that this occurs when mutation rates become roughly ten times higher than fragmentation rates (a-c). Evolutionary rescue is largely decreased in aggregated landscapes (d-f).

## DISCUSSION

Our study shows an increase of dispersal capacities in fragmented landscapes in the context of competition/colonization trade-offs. Aggregation acts as an opposite force, as decreased dispersal is selected in more aggregated landscapes. At high fragmentation and low aggregation, different strategies can be selected and can coexist, with better competitors in aggregated patches and better colonizers in isolated patches. Such an evolution of polymorphism allows a good global coverage of available space. When fragmentation increases with time, the rapid evolution of dispersal facilitates the survival of the metapopulation but this evolutionary rescue effect can only be observed in non-aggregated landscapes and when fragmentation rate is not too high.

The selection of more dispersive strategies in fragmented landscapes in a context of competition/colonization trade-offs is congruent with Tilman et al. (1994) which also relies on this trade-off. Other studies on the evolution of dispersal in spatially heterogeneous systems, but in the absence of a competition/colonization trade-off, show a reverse pattern, as dispersal is then counter selected because dispersal costs are enhanced by spatial heterogeneities (Hastings, 1983; Travis & Dytham, 1999). This highlights that patterns of selection strongly depend on the trade-off structure associated with dispersal traits. Recent works highlight the importance of dispersal syndromes (Raffard et al., 2022; Stevens et al., 2014), i.e. the fact that dispersal traits may be directly coupled to traits defining ecological interactions. The competition/colonization trade-off falls within this category, as variations of dispersal are directly coupled to competition hierarchies. Our results therefore highlight how such a syndrome could lead, for some landscapes, to the selection of higher dispersal, while works that consider evolution of dispersal alone (eg, Hastings, 1983; Travis and Dytham, 1999) would produce the reverse pattern. Application of either framework of course depends on the types of organisms that are considered and whether dispersal traits are competitively costly.

In particular, our model assumes that the fecundity of the organism under consideration, i.e. the number of offspring produced, increases with increasing dispersal distance. This provides an additional advantage for dispersive strategies that will produce many more offspring and therefore occupy space more quickly if there is no superior competitor present. Many previous models do not make this assumption and use constant fecundity (eg, Bonte et al., 2010; Travis and Dytham, 1999). This hypothesis may influence our results in several ways. With constant fecundity, selection of dispersive strategies is likely reduced, leading to a smaller mean dispersal in highly fragmented habitats, lower abundances in the landscape, and early extinctions. Dispersal polymorphism should stay present because isolated patches can only be reached by dispersive strategies. However, this assumption of increasing fecundity with increasing dispersal distance is not biologically irrelevant and can be linked to various groups of organisms. A certain amount of energy could be allocated in either a few large, poorly dispersing offspring or in many small, highly dispersing offspring. This rationale has been largely used in the context of variations of plant seed size (eg, Henery & Westoby, 2001; Muller-Landau et al., 2008). It could also apply in the context of social insect colonies. For instance, in ants with the production of large propagules consisting of a queen and workers that disperse over short distances or small propagules of a single queens dispersing over long distances (Cronin et al., 2013).

We explain the selection of higher dispersal in fragmented landscapes by two mechanisms. First, fragmentation decreases overall occupancy (on the entire grid). Thereby, when a patch is emptied, the number of possible colonizers (ie, of suitable filled patches) is reduced. This reduces the average competition level. The advantage of competitive strategies is therefore reduced. Second, fragmentation intensifies the strength of the competition for space which favors colonizers. Isolated patches in fragmented landscapes can only be exploited by highly dispersive strategies. Effects of fragmentation on mean levels of dispersal have led to contrasted results in empirical studies (Cheptou et al., 2017). Our results of a selection of higher dispersal is for instance coherent with empirical studies of nuthatches (*Sitta europaea*) in Belgium (Matthysen et al., 1995). Similarly, metapopulation study of the Glanville fritillary in Finland shows that isolated patches of the metapopulation act as positive filters for the more dispersive strategies (Hanski et al., 2004) and that this variation can be linked to allelic variations constraining flight metabolism (Haag et al., 2005). Conversely, a decrease of dispersal at higher fragmentation levels has been observed in various animal or plant species (Bonte et al., 2006; Cheptou et al., 2008; Schtickzelle et al., 2006). Spatial heterogeneity and competition decrease are two forces that act with opposite effects, the former decreasing dispersal (higher dispersal cost) while the latter increases it (decreased abundancies in fragmented landscapes, relax competition). The importance of each force varies among species and needs to be systematically considered to better predict changes in dispersal.

We found that aggregation largely reduces the selection of dispersal strategies, to the extent that such a selection cannot even be detected when aggregation is larger than 40%. This points out the importance of spatially explicit models. In a previous work, Hiebeler (2000) showed how mean field approximations provide accurate occupancy predictions for random fragmented landscapes, but not when aggregation exists. Similarly, we show here that while our results on dispersal evolution in random landscapes are coherent with mean field approximations (Tilman et al., 1994), such approximations do not provide qualitatively adequate variations when aggregation takes place. We explain the reduction of dispersal due to aggregation by the fact that it favors the replacement of colonizers by competitors because of a high probability to find a favorable patch next to another favorable patch. The landscape is there continuous, so that competition is selected in such localities. Such a result is in line with Bonte et al. (2010) who found an increase of local dispersal (our competitive strategy) and a decrease of long-distance dispersal (our colonizer strategy) in aggregated fragmentation scenarios. We therefore completely agree with the necessity of spatially explicit approaches to better understand the dynamics of fragmented metapopulations (Hiebeler, 2000). Here, a simple mean-field approach would yield an overestimation of dispersal evolution and of associated evolutionary rescue effects.

Our model is based on mutations and on the selection of certain phenotypes resulting from these mutations. While our results can be largely interpreted from a selection point of view, we explicitly account for stochastic components, both in the mutation process and in the random patch extinction process. This latter source of stochasticity leads to genetic drift in our simulations. Effects of drift are particularly visible for small metapopulation sizes (ie, on the brink of extinction), and indeed we observed broader distributions of phenotypic values in such conditions. To assess the importance of these stochastic components, we undertook 20 replications of each parameter combination in scenario one and 40 in scenario two. The consistent qualitative variations of dispersal distances however suggest a large role of selective processes.

We observe dispersal polymorphism when fragmentation is high and aggregation low to intermediate. Such landscapes contain a mix of large aggregates of patches and of isolated patches. The strategy favored in the aggregates of patches is more competitive, and the dispersal distance selected there depends on how loose the aggregates are. When aggregation produces continuous aggregates, the most competitive strategies are favored (dispersal distance close to 1, Fig. 3f), while when aggregates are looser, selected dispersal distances could be higher (Fig. 3f). In all cases, isolated patches favor more dispersive strategies. These results are coherent with previous theoretical studies that show how fragmentation can favor dispersal polymorphism. In particular, some of them showed that polymorphism is prevalent when few large patches (our patch aggregates) co-occur with small patches (our isolated patches) (Bonte et al., 2010; Massol et al., 2011; Parvinen, 2002; Parvinen et al., 2020). A large literature exists on how ecological dynamics of metapopulations under fragmentation leads to changes in persistence and to variations in diversity (Bascompte & Rodriguez, 2001; Bascompte & Solé, 1996; Ovaskainen et al., 2002; Solé et al., 2004). Previous works highlight the critical role of patchiness (Bascompte and Rodriguez, 2001) or of extinction thresholds (Bascompte and Solé, 1996). Here, our goal is rather to assess how fragmentation affects the evolution of dispersal and its eco-evolutionary consequences for the metapopulation dynamics. Such an evolution may in turn affect persistence (and diversity) when it fosters evolutionary rescue.

Evolutionary rescue can be construed as a race between speed of adaptation and of environmental deterioration. Hence, the faster the evolution and the slower the perturbation, the more likely the rescue. We observe that such expectations are met when fragmentation is random (no aggregation). Dispersal evolution delays the extinction of the population when fragmentation rate is low and mutation rate high. Fast selection of good dispersers then occurs. As these are adapted to occupy isolated patches, such strategies foster spatial rescue in the highly fragmented landscapes. Slow evolution would not allow that. At the onset of fragmentation, the grid is continuous, and mostly occupied by competitors. If fragmentation is too fast, there is no time for dispersers to appear and become selected and to fill isolated patches. Interestingly, our study shows that rescue happens as a jump phenomenon, being only possible when mutation rates are higher (10 times higher in our model) than perturbation rates. No evolutionary rescue occurs when fragmentation is aggregated. Aggregation delays extinction by itself even without evolution. Under high fragmentation and aggregation levels, suitable patches make small continuous groups that facilitate the local persistence of competitors. In an aggregated context, dispersal evolution is absent or strongly constrained (blue curves on Fig. 2a) so that little evolutionary potential exists. In the absence of such an adaptive potential, evolutionary rescue cannot act. Our results on the possibility of rescue through evolutionary changes of dispersal agree with former theoretical works where fragmentation either stemmed from climatic changes (Boeye et al., 2012) or from heterogeneities in mortality (Heino & Hanski, 2001). While in the actual context of fast environmental changes, it may seem complicated for species to evolve quickly enough (10 times faster than the perturbation), several examples of fast evolution of dispersal in fragmented systems have been reported (reviewed in Cheptou et al., 2017). Whether such evolution are sufficient to affect long term metapopulation persistence is however unknown. The fact that evolutionary rescue does not happen here in aggregated landscapes also has important implications. The current fragmentation of habitats is a complex non-random process that may be frequently auto-correlated in space, therefore producing aggregated structures. For instance, the construction of additional urban areas next to existing urban areas creates aggregated landscapes. Studies from the Tabriz Metropolitan Area (Iran) show that the destruction of suitable habitats surrounding the cities result in the creation of aggregated non-suitable patches (Dadashpoor et al., 2019b, 2019a). Aggregation of fragmentation can also be linked to the displacement and development of agricultural activities. In Beijing City, China, landscape patterns show important and complicated changes in the distribution of urban and agricultural lands. Economic development expands cultivated land and construction into forests and grasslands resulting in aggregated and less diverse landscapes (Li et al., 2017). We propose that when fragmentation happens in such aggregated ways, evolution will likely play a minor role in the maintenance of the metapopulation.

Our study highlights the importance of considering dispersal syndromes (here through the competition/colonization trade-off) and the structuration of habitat fragmentation to better understand how dispersal evolves in disturbed landscapes. We acknowledge that our model is quite simple and can only be used to provide baseline scenarios. For instance, fragmentation can create changes not only in competition intensity, but also in other community aspects (eg, presence of mutualists and enemies, see Cheptou et al., 2017). While we simply focus on the colonization-competition trade-off, evolutionary changes can also involve other phenotypic traits. Colonization of empty patches, usually free of conspecifics, could for instance lead to the fast evolution of intrinsic growth rates (Williams et al., 2019). We hope that the results we present here will motivate efforts to better understand the multidimensionality of dispersal evolution and its implications for the future of biodiversity.

## Supporting information

Supplementaty informations

## DATA AVAIBILITY

Model and analysis scripts are available on github: https://github.com/bfinand/Model_dispersal_evolution

## REFERENCES

Bascompte, J., & Rodriguez, M. A. (2001). Habitat patchiness and plant species richness. Ecology Letters, 4(5), 417–420. https://doi.org/10.1046/j.1461-0248.2001.00242.x

Bascompte, J., & Solé, R. V. (1996). Habitat fragmentation and extinction thresholds in spatially explicit models. Journal of Animal Ecology, 65, 465–473.

Bell, G. (2017). Evolutionary Rescue. Annual Review of Ecology, Evolution, and Systematics, 48(1), 605–627. https://doi.org/10.1146/annurev-ecolsys-110316-023011

Boeye, J., Travis, J. M. J., Stoks, R., & Bonte, D. (2012). More rapid climate change promotes evolutionary rescue through selection for increased dispersal distance. Evolutionary Applications, 6(2), 353–364. https://doi.org/10.1111/eva.12004

Bonte, D., Borre, J. Vanden, Lens, L., & Jean-Pierre Maelfait. (2006). Geographical variation in wolf spider dispersal behaviour is related to landscape structure. Animal Behaviour, 72(3), 655–662. https://doi.org/10.1016/j.anbehav.2005.11.026

Bonte, D., Hovestadt, T., & Poethke, H. J. (2010). Evolution of dispersal polymorphism and local adaptation of dispersal distance in spatially structured landscapes. Oikos, 119, 560–566. https://doi.org/10.1111/j.1600-0706.2009.17943.x

Calcagno, V., Mouquet, N., Jarne, P., & David, P. (2006). Coexistence in a metacommunity: The competition-colonization trade-off is not dead. Ecology Letters, 9, 897–907. https://doi.org/10.1111/j.1461-0248.2006.00930.x

Carlson, S. M., Cunningham, C. J., & Westley, P. A. H. (2014). Evolutionary rescue in a changing world. Trends in Ecology & Evolution, 29(9), 521–530. https://doi.org/10.1016/j.tree.2014.06.005

Charlesworth, D., & Charlesworth, B. (1987). Inbreeding depression and its evolutionary consequences. Annual Review of Ecology and Systematics, 18(1), 237–268. https://doi.org/10.1146/annurev.es.18.110187.001321

Cheptou, P.-O., Carrue, O., Rouifed, S., & Cantarel, A. (2008). Rapid evolution of seed dispersal in an urban environment in the weed Crepis sancta. Proceedings of the National Academy of Sciences, 105(10), 3796–3799. https://doi.org/10.1073/pnas.0708446105

Cheptou, P.-O., Hargreaves, A. L., Bonte, D., & Jacquemyn, H. (2017). Adaptation to fragmentation: evolutionary dynamics driven by human influences. Philosophical Transactions of the Royal Society B: Biological Sciences, 372(1712), 20160037. https://doi.org/10.1098/rstb.2016.0037

Cote, J., Bestion, E., Jacob, S., Travis, J., Legrand, D., & Baguette, M. (2017). Evolution of dispersal strategies and dispersal syndromes in fragmented landscapes. Ecography, 40, 56–73. https://doi.org/10.1111/ecog.02538

Cronin, A. L., Loeuille, N., & Monnin, T. (2016). Strategies of offspring investment and dispersal in a spatially structured environment: a theoretical study using ants. BMC Ecology, 16(1), 4. https://doi.org/10.1186/s12898-016-0058-z

Cronin, A. L., Molet, M., Doums, C., Monnin, T., & Peeters, C. (2013). Recurrent Evolution of Dependent Colony Foundation Across Eusocial Insects. Annual Review of Entomology, 58(1), 37–55. https://doi.org/10.1146/annurev-ento-120811-153643

Dadashpoor, H., Azizi, P., & Moghadasi, M. (2019a). Analyzing spatial patterns, driving forces and predicting future growth scenarios for supporting sustainable urban growth: Evidence from Tabriz metropolitan area, Iran. Sustainable Cities and Society, 47, 101502. https://doi.org/10.1016/j.scs.2019.101502

Dadashpoor, H., Azizi, P., & Moghadasi, M. (2019b). Land use change, urbanization, and change in landscape pattern in a metropolitan area. Science of The Total Environment, 655, 707–719. https://doi.org/10.1016/j.scitotenv.2018.11.267

Duputié, A., & Massol, F. (2013). An empiricist’s guide to theoretical predictions on the evolution of dispersal. Interface Focus, 3(6), 20130028. https://doi.org/10.1098/rsfs.2013.0028

Fahrig, L. (2003). Effects of Habitat Fragmentation on Biodiversity. Annual Review of Ecology, Evolution, and Systematics, 34, 487–515. https://doi.org/10.1146/annurev.ecolsys.34.011802.132419

Fronhofer, E. A., Stelz, J. M., Lutz, E., Poethke, H. J., & Bonte, D. (2014). Spatially correlated extinctions select for less emigration but larger dispersal distances in the spider mite Tetranychus urticae. Evolution, 68(6), 1838–1844. https://doi.org/10.1111/evo.12339

Gandon, S. (1999). Kin Competition, the Cost of Inbreeding and the Evolution of Dispersal. Journal of Theoretical Biology, 200(4), 345–364. https://doi.org/10.1006/jtbi.1999.0994

Geritz, S. A. H., van der Meijden, E., & Metz, J. A. J. (1999). Evolutionary Dynamics of Seed Size and Seedling Competitive Ability. Theoretical Population Biology, 55(3), 324–343. https://doi.org/10.1006/tpbi.1998.1409

Gomulkiewicz, R., & Holt, R. D. (1995). When does Evolution by Natural Selection Prevent Extinction ? Evolution, 49(1), 201–207.

Gyllenberg, M., Parvinen, K., & Dieckmann, U. (2002). Evolutionary suicide and evolution of dispersal in structured metapopulations. Journal of Mathematical Biology, 45(2), 79–105. https://doi.org/10.1007/s002850200151

Haag, C. R., Saastamoinen, M., Marden, J. H., & Hanski, I. (2005). A candidate locus for variation in dispersal rate in a butterfly metapopulation. Proceedings of the Royal Society B: Biological Sciences, 272(1580), 2449–2456. https://doi.org/10.1098/rspb.2005.3235

Hamilton, W. D., & May, R. M. (1977). Dispersal in stable habitats. Nature, 269(5629), 578–581. https://doi.org/10.1038/269578a0

Hanski, I., Erälahti, C., Kankare, M., Ovaskainen, O., & Sirén, H. (2004). Variation in migration propensity among individuals maintained by landscape structure. Ecology Letters, 7(10), 958–966. https://doi.org/10.1111/j.1461-0248.2004.00654.x

Hastings, A. (1983). Can spatial variation alone lead to selection for dispersal? Theoretical Population Biology, 24, 244–251. https://doi.org/10.1016/0040-5809(83)90027-8

Heino, M., & Hanski, I. (2001). Evolution of Migration Rate in a Spatially Realistic Metapopulation Model. The American Naturalist, 157(5), 495–511. https://doi.org/10.1086/319927

Henery, M. L., & Westoby, M. (2001). Seed mass and seed nutrient content as predictors of seed output variation between species. Oikos, 92(3), 479–490. https://doi.org/10.1034/j.1600-0706.2001.920309.x

Hiebeler, D. (2000). Populations on fragmented landscapes with spatially structured heterogeneities: landscape generation and local dispersal. Ecology, 81(6), 1629–1641.

Levins, R. (1969). Some Demographic and Genetic Consequences of Environmental Heterogeneity for Biological Control. Bulletin of the Entomological Society of America, 15(3), 237–240. https://doi.org/10.1093/besa/15.3.237

Li, H., Peng, J., Yanxu, L., & Yi’na, H. (2017). Urbanization impact on landscape patterns in Beijing City, China: A spatial heterogeneity perspective. Ecological Indicators, 82, 50–60. https://doi.org/10.1016/j.ecolind.2017.06.032

Massol, F., Duputié, A., David, P., & Jarne, P. (2011). Asymmetric patch size distribution leads to disruptive selection on dispersal. Evolution, 65(2), 490–500. https://doi.org/10.1111/j.1558-5646.2010.01143.x

Matthysen, E., Adriaensen, F., Dhondt, A. A., & Dhondt, A. A. (1995). Dispersal Distances of Nuthatches, Sitta europaea, in a Highly Fragmented Forest Habitat. Oikos, 72(3), 375. https://doi.org/10.2307/3546123

Muller-Landau, H. C., Wright, S. J., Calderón, O., Condit, R., & Hubbell, S. P. (2008). Interspecific variation in primary seed dispersal in a tropical forest. Journal of Ecology, 96(4), 653–667. https://doi.org/10.1111/j.1365-2745.2008.01399.x

Oldfather, M. F., Van Den Elzen, C. L., Heffernan, P. M., & Emery, N. C. (2021). Dispersal evolution in temporally variable environments: implications for plant range dynamics. American Journal of Botany, 108(9), 1584–1594. https://doi.org/10.1002/ajb2.1739

Ovaskainen, O., Sato, K., Bascompte, J., & Hanski, I. (2002). Metapopulation Models for Extinction Threshold in Spatially Correlated Landscapes. Journal of Theoretical Biology, 215(1), 95–108. https://doi.org/10.1006/jtbi.2001.2502

Parvinen, K. (2002). Evolutionary branching of dispersal strategies in structured metapopulations. Journal of Mathematical Biology, 45(2), 106–124. https://doi.org/10.1007/s002850200150

Parvinen, K., Ohtsuki, H., & Wakano, J. Y. (2020). Evolution of dispersal in a spatially heterogeneous population with finite patch sizes. Proceedings of the National Academy of Sciences, 117(13), 7290–7295. https://doi.org/10.1073/pnas.1915881117

Phillips, B. L., Brown, G. P., Webb, J. K., & Shine, R. (2006). Invasion and the evolution of speed in toads. Nature, 439(7078), 803–803. https://doi.org/10.1038/439803a

Pulliam, H. R. (1988). Sources, Sinks, and Population Regulation. The American Naturalist, 132(5), 652–661. https://doi.org/10.1086/284880

Raffard, A., Bestion, E., Cote, J., Haegeman, B., Schtickzelle, N., & Jacob, S. (2022). Dispersal syndromes can link intraspecific trait variability and meta-ecosystem functioning. Trends in Ecology & Evolution, 37(4), 322–331. https://doi.org/10.1016/j.tree.2021.12.001

Ronce, O. (2007). How does it feel to be like a rolling stone? Ten questions about dispersal evolution. Annual Review of Ecology, Evolution, and Systematics, 38(1), 231–253. https://doi.org/10.1146/annurev.ecolsys.38.091206.095611

Schtickzelle, N., Mennechez, G., & Baguette, M. (2006). Dispersal depression with habitat fragmentation in the bog fritillary butterfly. Ecology, 87(4), 1057–1065. https://doi.org/10.1890/0012-9658(2006)87[1057:DDWHFI]2.0.CO;2

Sciaini, M., Fritsch, M., Scherer, C., & Simpkins, C. E. (2018). NLMR and landscapetools: An integrated environment for simulating and modifying neutral landscape models in R. Methods in Ecology and Evolution, 9(11), 2240–2248. https://doi.org/10.1111/2041-210X.13076

Smith, C. C., & Fretwell, S. D. (1974). The Optimal Balance between Size and Number of Offspring. The American Naturalist, 108(962), 499–506. https://doi.org/10.1086/282929

Solé, R. V., Alonso, D., & Saldaña, J. (2004). Habitat fragmentation and biodiversity collapse in neutral communities. Ecological Complexity, 1(1), 65–75. https://doi.org/10.1016/j.ecocom.2003.12.003

Stevens, V. M., Whitmee, S., Le Galliard, J.-F., Clobert, J., Böhning-Gaese, K., Bonte, D., Brändle, M., Matthias Dehling, D., Hof, C., Trochet, A., & Baguette, M. (2014). A comparative analysis of dispersal syndromes in terrestrial and semi-terrestrial animals. Ecology Letters, 17(8), 1039–1052. https://doi.org/10.1111/ele.12303

Tilman, D. (1994). Competition and Biodiversity in Spatially Structured Habitats. Ecology, 75(1), 2–16. https://doi.org/10.2307/1939377

Tilman, D., Lehman, C. L., & Yin, C. (1997). Habitat Destruction, Dispersal, and Deterministic Extinction in Competitive Communities. The American Naturalist, 149(3), 407–435. https://doi.org/10.1086/285998

Tilman, D., May, R. M., Lehman, C. L., & Nowak, M. A. (1994). Habitat destruction and the extinction debt. Nature, 371(6492), 65–66. https://doi.org/10.1038/371065a0

Travis, J. M. J., Delgado, M., Bocedi, G., Baguette, M., Bartoń, K., Bonte, D., Boulangeat, I., Hodgson, J. A., Kubisch, A., Penteriani, V., Saastamoinen, M., Stevens, V. M., & Bullock, J. M. (2013). Dispersal and species’ responses to climate change. Oikos, 122, 1532–1540. https://doi.org/10.1111/j.1600-0706.2013.00399.x

Travis, J. M. J., & Dytham, C. (1999). Habitat persistence, habitat availability and the evolution of dispersal. Proceedings of the Royal Society B: Biological Sciences, 266, 723–728. https://doi.org/10.1098/rspb.1999.0696

Tung, S., Mishra, A., Shreenidhi, P. M., Sadiq, M. A., Joshi, S., Sruti, V. R. S., & Dey, S. (2018). Simultaneous evolution of multiple dispersal components and kernel. Oikos, 127(1), 34–44. https://doi.org/10.1111/oik.04618

Williams, J. L., Hufbauer, R. A., & Miller, T. E. X. (2019). How Evolution Modifies the Variability of Range Expansion. Trends in Ecology & Evolution, 34(10), 903–913. https://doi.org/10.1016/j.tree.2019.05.012

Yu, D. W., & Wilson, H. B. (2001). The Competition-Colonization Trade-off Is Dead; Long Live the Competition-Colonization Trade-off. The American Naturalist, 158(1), 49–63. https://doi.org/10.1086/320865

