## Supplementaty informations for "Evolution of dispersal and the maintenance of fragmented metapopulations"

**SUPPLEMENTARY MATERIALS**

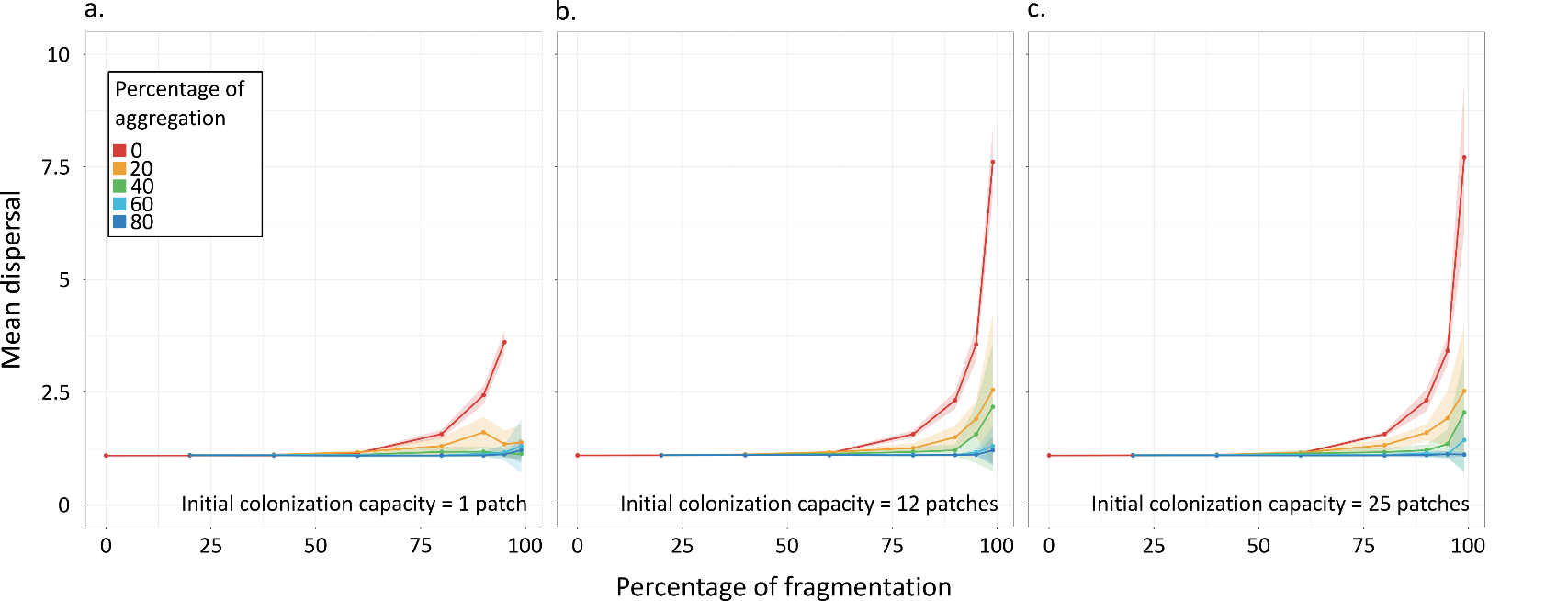

**Figure S1** : Dispersal (mean +/- SD) at the end of simulations (5 000 time steps) versus environment fragmentation, aggregation and initial colonization capacity. We started the simulation with 1 individual in the grid and with an initial colonization capacity of a) 1 patch b) 12 patches c) 25 patches. All simulations stop quickly for a percentage of fragmentation of 99%, a percentage of aggregation of 0% and an initial colonization capacity of 1 patch that explain the absence of data in a) red line. We observed that the initial colonization capacity and the initial number of populations on the grid do not change the outcomes of the model.

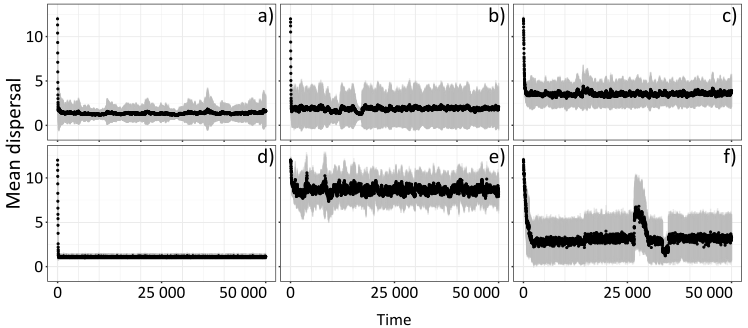

**Figure S2:** Mean dispersal in the grid over time for different examples of simulation. Black dots are the mean of the trait distribution for each time step and grey area show the standard deviation. a) percentage of fragmentation of 80 and aggregation of 20 b) percentage of fragmentation of 90 and aggregation of 20 c) percentage of fragmentation of 95 and aggregation of 0 d) percentage of fragmentation of 95 and aggregation of 60 e) percentage of fragmentation of 99 and aggregation of 0 f) percentage of fragmentation of 99 and aggregation of 20

These plots highlight that despite important stochastic components, our model evolves toward a stable distribution much faster than the simulation time we here consider. Note that plots vary largely in their parameter conditions and are therefore representative of our set of simulations. The stability was checked visually for all simulations. While stochastic components make the use of an exact threshold difficult to determine stationarity, they also provide important insights regarding the relative roles of selection and drift, two key processes which have implications in the maintenance of metapopulations.
